# Resting-state functional MRI derivatives: A dataset derived from the The Comprehensive Assessment of Neurodegeneration and Dementia Study

**DOI:** 10.1101/2025.10.14.682410

**Authors:** Natasha Clarke, Hao-Ting Wang, Désirée Lussier, Arnaud Boré, Loic Tetrel, Camille Beaudoin, Samir Das, Randi Pilon, Alan C. Evans, Howard Chertkow, Roger A. Dixon, AmanPreet Badhwar, Simon Duchesne, Lune Bellec

**Affiliations:** Centre de Recherche de l’Institut Universitaire de Gériatrie de Montréal (CRIUGM), 4565 chemin Queen Mary, Montréal, QC, H3W 1W5, Canada; Département de Psychologie, Université de Montréal; QC H2V 2S9, Montréal, Canada; Montreal Neurological Institute and Hospital, McGill University; QC H3A 2B4, Montreal, Canada; Lady Davis Institute for Medical Research, Jewish General Hospital; QC H3T 1E2, Montreal, Canada; Baycrest Institute for Research and Education and Department of Medicine, University of Toronto; Department of Neurology and Neurosurgery, McGill University, Montreal, Canada; Department of Psychology (Science) and Neuroscience and Mental Health Institute, University of Alberta, Edmonton, Alberta T6G 2E9; Multiomics Investigation of Neurodegenerative Diseases (MIND) lab, 4545 Queen Mary Road, Montréal, QC, H3W 1W6, Canada; Department of Pharmacology and Physiology, Faculty of Medicine, Université de Montréal, 2900 boulevard Édouard-Montpetit, Montréal, QC, H3T 1J4, Canada; Institute of Biomedical Engineering, Université de Montréal, 2960 chemin de la Tour, Montréal, QC, H3T 1J4, Canada; Quebec Heart and Lung Research Institute, Quebec City, Canada; Department of radiology and nuclear medicine, Université Laval, Quebec City, Canada

## Abstract

Resting-state functional connectivity (RSFC) holds promise for the detection and characterisation of dementia. The Comprehensive Assessment of Neurodegeneration and Dementia (COMPASS-ND) Study, by the Canadian Consortium on Neurodegeneration in Aging (CCNA), provides a unique resource to study deeply phenotyped neurodegenerative conditions. We present RSFC derivatives for 784 participants (data release 7 of the cohort) who were either cognitively unimpaired or diagnosed primarily with Alzheimer’s disease (AD), mixed dementia (AD with a vascular component), mild cognitive impairment (MCI), vascular MCI, frontotemporal dementia, Parkinson’s disease with or without MCI or dementia, Lewy body disease or subjective cognitive impairment. Functional MRI scans were preprocessed using fMRIPrep, and time-series and whole-brain connectomes generated using three atlases at multiple resolutions, denoised using seven different techniques. High-motion artifacts were managed using a liberal quality control threshold appropriate for an older clinical population, resulting in data from 680 participants. These derivatives are made available to the research community to accelerate research on RSFC biomarkers of neurodegenerative disease, reducing duplication of effort, saving computational resources, and improving standardisation across studies.

## Background and summary

Dementia affects over 57 million people worldwide^1^, and over 650,000 Canadians^2^, with a devastating impact on health and quality of life. Alzheimer’s disease (AD) is the leading cause, followed by vascular dementia^3^, while other less common forms include frontotemporal dementia (FTD), Parkinson’s disease (PD), and Lewy body disease (LBD), each with distinct neurodegeneration patterns and symptom profiles. Early and accurate diagnosis, including identifying those at an increased risk of progression to dementia such as mild cognitive impairment (MCI) and subjective cognitive impairment (SCI)^4,5^, is crucial, as pathology starts to accumulate years prior to overt symptoms. The identification of biomarkers is a core goal of neuroscience research that can help with early detection, risk stratification and disease monitoring.

Resting-state functional connectivity (RSFC) is a promising biomarker for detecting early brain changes associated with dementia. RSFC, derived from the strength of correlation between functional MRI (fMRI) time-series whilst the brain is at rest, captures synchronous fluctuations in the blood oxygenation level dependent (BOLD) signals between brain regions^6^. Regions with highly synchronous BOLD signals are considered functionally related, and can be studied as a whole-brain connectivity matrix (a connectome), or grouped into networks representing the brain’s intrinsic architecture. Commonly studied networks include the default mode, somatomotor, dorsal attention, visual, ventral attention, limbic and frontoparietal^7^. Evidence suggests resting-state networks are disrupted in neurodegenerative conditions such as AD^8,9^, PD^10^, and vascular dementia^11^. Perturbations are also observed in MCI^12^, vascular-MCI^13^, and preclinical populations at increased risk of AD, including cognitively unimpaired individuals with amyloid-beta plaques^14^ or a family history of AD^15^. Resting-state fMRI is also non-invasive and does not require engagement with a task, which is beneficial for neurodegenerative populations. However, the development of effective RSFC biomarkers is hampered by disease heterogeneity, symptom overlap and a lack of longitudinal data.

The Comprehensive Assessment of Neurodegeneration and Dementia (COMPASS-ND) Study is a pan-Canadian observational, longitudinal study of age-related dementia, and the signature cohort of the Canadian Consortium on Neurodegeneration in Aging (CCNA). The study is unique in its breadth and depth, including cognitively unimpaired individuals alongside those with cognitive impairment and a range of neurodegenerative profiles, including mixed pathologies. To date, 1,173 participants have completed the baseline assessment (Time 1), with another 400 participants to be recruited over the coming years. Longitudinal follow-up includes an abbreviated Time 2 assessment, the timing of which was disrupted by the COVID-19

pandemic, resulting in visits occurring on average 3 years post-baseline. A full-protocol Time 3 assessment, which includes repeat neuroimaging, was recently initiated, and will occur 6-8 years post-baseline. Participants are recruited from 30 specialty sites across Canada and undergo deep phenotyping, clinical and cognitive assessments, genetic and biospecimen analysis, optional brain donation, and must consent to brain imaging^16^. MRI acquisitions follow a harmonized protocol^17^ designed to minimize inter-manufacturer variations - a major challenge for multi-site studies^18,19^. The dataset provides researchers with an unprecedented opportunity to study multiple age-related dementias within the same cohort, improving understanding of heterogeneous dementias and aiding differential diagnosis. In this paper, we describe the preprocessing of fMRI data and the extraction of time-series and whole-brain functional connectome derivatives, essential for studying connectivity alterations in neurodegeneration and moving towards clinical adoption of RSFC as a biomarker.

## Methods

### Participants

Participants aged 50-90 were enrolled in the COMPASS-ND cohort (registered on clinicaltrials.gov: NCT03402919). They were either cognitively unimpaired (CU) older adults or diagnosed clinically with subjective cognitive impairment (SCI); mild cognitive impairment (MCI); vascular-MCI (V-MCI); Alzheimer’s disease (AD); mixed dementia (AD and a vascular component); Lewy body disease (LBD); Parkinson’s disease either with MCI (PD-MCI), no cognitive impairment (PD), or dementia (PDD); or frontotemporal dementia (FTD), which included behavioural variant FTD, primary progressive aphasia (PPA), cortico-basal syndrome (CBS) and progressive supranuclear palsy (PSP). Diagnoses were made by investigators at each site based on the following published criteria: SCI^20^, MCI^21^, V-MCI^3,22^, AD or mixed dementia^23^, LBD^24^, PD^25–28^, behavioral variant FTD^29^, PPA^30,31^, CBS^32^, PSP^33^. Ethical approval was obtained from institutional review boards at each site, and all participants gave informed consent. For detailed information on the protocol, including inclusion/exclusion criteria, see Chertkow and colleagues^16^. Here we focus on baseline data release 7, which includes resting-state fMRI for 784 participants.

### Data acquisition

Participants completed 3T MRI following the Canadian Dementia Imaging Protocol (CDIP; www.cdip-pcid.ca). CDIP is a standardized, tri-vendor (GE Healthcare, Philips Medical and Siemens Medical Systems) acquisition protocol that can be completed in approximately 45 minutes^17^. It includes 3D T1-weighted, T2/PD-weighted, fluid-attenuated inversion recovery (FLAIR), T2* and diffusion-weighted sequences, and a task-free, eyes-open resting state blood-oxygen-level-dependent (BOLD) acquisition. Functional T2*-weighted images were acquired using a BOLD-sensitive single-shot echo-planar (EPI) sequence at 3.5 x 3.5 x 3.5 mm^3^ resolution, with a TR of 2110 msec (GE: 2500 msec) and 300 volumes over approximately 11 minutes. For a complete description of the CDIP protocol see Duchesne and colleagues^17^. Exam cards and other technical details for GE Discovery, Philips Achieva Ingenia, and Siemens Magnetom TRIO and PRISMA scanners are available on the CDIP website (www.cdip-pcid.ca).

### MRI preprocessing

DICOM images were obtained from the MRI scanner and converted to NIfTI format using dcm2bids, organized according to the Brain Imaging Data Structure (BIDS) standard^34^. Preprocessing was performed using fMRIPrep 20.2.7 long-term support (LTS) branch^35^, which is based on Nipype 1.7.0^36,37^. The fMRIprep pipeline performs basic processing steps including coregistration, normalization, unwarping, noise component extraction, segmentation and skullstripping, using well-known software packages including FSL, ANTs, FreeSurfer and AFNI. Many internal operations leverage Nilearn 0.6.2, primarily within the functional workflow. More information on the pipeline is available at https://fmriprep.org/en/stable/index.html, and full details of the anatomical and functional outputs are provided in the supplementary materials.

#### Quality control

Automatic quality inspection of anatomical and functional data was performed using the Giga AutoQC tool v0.3.3, a BIDS app developed in the lab and available on GitHub (https://github.com/SIMEXP/giga_auto_qc). Participant head motion was assessed using mean framewise displacement (FD) before and after scrubbing. FD, calculated from six motion parameters per participant, indexes movement of the head between volumes, and is implemented in fMRIPrep following the formula proposed by Power and colleagues^38^. The mean FD across all time points was calculated. To ensure data quality and reduce motion artifacts, frames exceeding a threshold were removed (scrubbed). A conservative threshold of 0.2mm FD is common for determining low-motion, high quality data in healthy populations^39^. However, due to high motion in the COMPASS-ND dataset, a threshold of 0.2mm resulted in the loss of a substantial number of frames. We therefore applied a more liberal threshold of 0.55mm, sufficient to reduce the impact of motion while preserving more timeframes^40^. The proportion of volumes retained after scrubbing was calculated, with a threshold set at 0.5, meaning that a subject should have at least 150 low motion volumes (approximately 5 minutes of data). The Sørensen–Dice coefficient was computed between the functional mask and a group fMRI mask evaluated with NiLearn^41^, and between the anatomical mask resampled to the MNI template and the corresponding MNI template^42^. Thresholds were set at .99 for anatomical and .89 for functional Dice. A subject failed QC if they did not meet the anatomical threshold, or failed any functional thresholds.

#### Time-series and connectome generation

Time-series extraction and connectome generation were performed using the Giga Connectome BIDS app v0.4.1^43^ developed in the lab, based on Python and Nilearn. Time-series were extracted from regions of interest (ROIs) defined by three different atlases at multiple resolutions (in parentheses): Multiresolution Intrinsic Segmentation Template (MIST)^44^ (7, 12, 20, 36, 64, 122, 197, 325, 444), Schaefer 7^45^ (100, 200, 300, 400, 500, 600, 800) and Dictionary of Functional Modes (DiFuMo)^46^ (114, 200, 372, 637, 1158), and averaged per ROI. Whole-brain connectomes (connectivity matrices) were computed at the subject level using Pearson’s correlation between ROI time-series, and a group-level connectome was obtained by averaging subject-level connectomes. Seven confound regression strategies were implemented to reduce noise. A simple strategy based on Fox and colleagues^47^ regressed motion parameters and tissue signal. Two scrubbing strategies based on Power and colleagues^38^ excluded time points with motion above the liberal (0.55mm) or strict (0.2mm) framewise displacement threshold, while also regressing motion parameters and tissue signal. Finally, a component based strategy (CompCor)^48^ that uses signal from regions deemed likely to be noise (e.g. white matter) to model grey matter noise was implemented. All strategies were implemented with and without global signal regression, except CompCor. For an in-depth comparison of these strategies using different data see Wang and colleagues^40^. Time-series and confound regressors from fMRIPrep are also provided, enabling users to apply their preferred atlas and denoising strategies.

### Data records

#### Data access

Access to the RSFC derivatives described here is made accessible to CCNA members and collaborators upon request. For information on CCNA membership and data access see https://ccna-ccnv.ca/membership/ and https://ccna-ccnv.ca/publication-policy/ and https://ccna-ccnv.ca/publication-policy/. A data usage agreement is available via the CCNA LORIS repository (https://ccna.loris.ca/), a web-based platform for neuroimaging data and project management^49,50^. Following approval, users can download derivatives from the CCNA ProFTP server.

#### Data organisation

The dataset is organised according to the BIDS format, a community-driven standard that improves efficiency, reproducibility and interoperability of research data^34^. A participants.tsv file lists the subjects included in the release, with basic phenotype information (age at scan, sex, site of MRI data collection, handedness and years of education), while a metadata.json file provides additional details. Four main folders are available for download, depending on the user’s analysis needs:

1. *compass-nd_fmriprep-20.2.7lts/bids_release_7/fmriprep-20.2.7lts*: Contains subject-and session-specific preprocessed anatomical (*anat*) and functional (*func*) MRI data. Functional directories (*func)* include preprocessed BOLD time-series (*\*_desc-preproc_bold.nii.gz*), confound regressors (*\*_desc-confounds_timeseries.tsv*), and brain masks. Anatomical directories (*anat)* include T1-weighted images (*\*_desc-preproc_T1w.nii.gz*), tissue probability maps, and cortical surface files (e.g., *\*_hemi-L_pial.surf.gii*). Freesurfer outputs are stored under *sourcedata/freesurfer/sub-<ID>*.
2. *compass-nd_connectome-0.4.1:* Contains subject- and session-level functional derivatives stored in HDF5 format. Data were derived using multiple atlases and denoising strategies, with files named *sub-<ID>_atlas-<ATLAS>_desc-<STRATEGY>.h5*, containing time-series and connectomes. Timeseries are of shape [n_regions, n_timepoints] and named \**atlas-<ATLAS>_desc-<RESOLUTION>_timeseries*. Connectomes are of shape [n_regions, n_regions] and named \**atlas-<ATLAS>_desc-<RESOLUTION>_connectome*. For DiFuMo and MIST atlases, the resolution indicates the number of dimensions; for the Schaefer atlas it indicates the number of parcels and networks. Group-level atlases and a grey matter mask are stored in *working_directory/groupmasks/tpl-MNI152NLin2009cAsym*.
3. *compass-nd_giga_auto_qc-0.3.3:* Automated QC reports per subject using a strict scrubbing threshold of .2mm, named *task-rest_report.tsv*.
4. *Compass-nd_giga_auto_qc-0.3.3_scrub.5*: Automated QC reports per subject using a liberal scrubbing threshold of .55mm, also named *task-rest_report.tsv*.

### Technical validation

A total of 788 MRI scans from 784 participants were preprocessed and underwent automatic QC based on motion and alignment (see figure 1 for flowchart). 783 passed anatomical QC and 690 passed functional QC using a liberal head motion threshold of framewise displacement (<.55mm). 683 scans passed both anatomical and functional QC, with the anatomical failures due to anatomical Dice coefficients marginally below the threshold. Time-series extraction failed for three participants, resulting in connectome and time-series data from 680 scans collected from 680 participants. These participants included 11 different diagnostic categories, according to the diagnosis under “Latest CND Dx” in central CCNA data. Users may wish to use alternative diagnostic data depending on their study objectives. In order of the number of individual connectomes/time-series data available (in parentheses), these include MCI (183), SCI (109), V-MCI (98), CU (78), PD (56), AD (45), mixed (45), PD-MCI (24), FTD (15), LBD (16) and PDD (11). Summary demographics are provided in Table 1.

**Table 1.** Sample size and demographic information for the 680 participants with scans that passed anatomical and functional automatic QC, and available derivatives. Continuous variables (age, education, MoCA) are reported as mean ± SD and were compared across groups using the Kruskal-Wallis test. Categorical variables (sex) are reported as n (%) and were compared using the chi-square test.

| **Diagnosis** | **N female** | **Age** | **Education** | **MoCA** |
| --- | --- | --- | --- | --- |
| MCI (n=183) | 72 (39.3%) | 72.1 (6.9) | 15.6 (3.9) | 23.7 (3.0) |
| SCI (n=109) | 79 (72.5%) | 69.7 (5.7) | 16.4 (3.5) | 27.4 (1.8) |
| V-MCI (n=98) | 45 (45.9%) | 75.6 (5.9) | 15.1 (3.6) | 23.1 (3.4) |
| CU (n=78) | 61 (78.2%) | 69.9 (5.5) | 15.9 (3.5) | 27.8 (1.6) |
| PD (n=56) | 24 (42.9%) | 66.8 (7.2) | 16.0 (3.6) | 27.6 (1.8) |
| AD (n=45) | 16 (35.6%) | 72.7 (6.8) | 15.3 (4.2) | 18.4 (4.0) |
| Mixed (n=45) | 21 (46.7%) | 79.2 (6.3) | 15.0 (3.5) | 18.5 (3.5) |
| PD-MCI (n=24) | 6 (25.0%) | 70.0 (7.7) | 15.8 (3.3) | 21.8 (4.1) |
| LBD (n=16) | 2 (12.5%) | 71.2 (9.1) | 13.7 (3.6) | 18.6 (3.5) |
| FTD (n=15) | 9 (60.0%) | 64.8 (8.7) | 15.1 (4.2) | 19.3 (4.8) |
| PDD (n=11) | 1 (9.1%) | 76.6 (6.2) | 16.8 (4.5) | 19.6 (4.2) |
| *p*-value | <0.001 | <0.001 | 0.103 | <0.001 |

**Figure 1.**
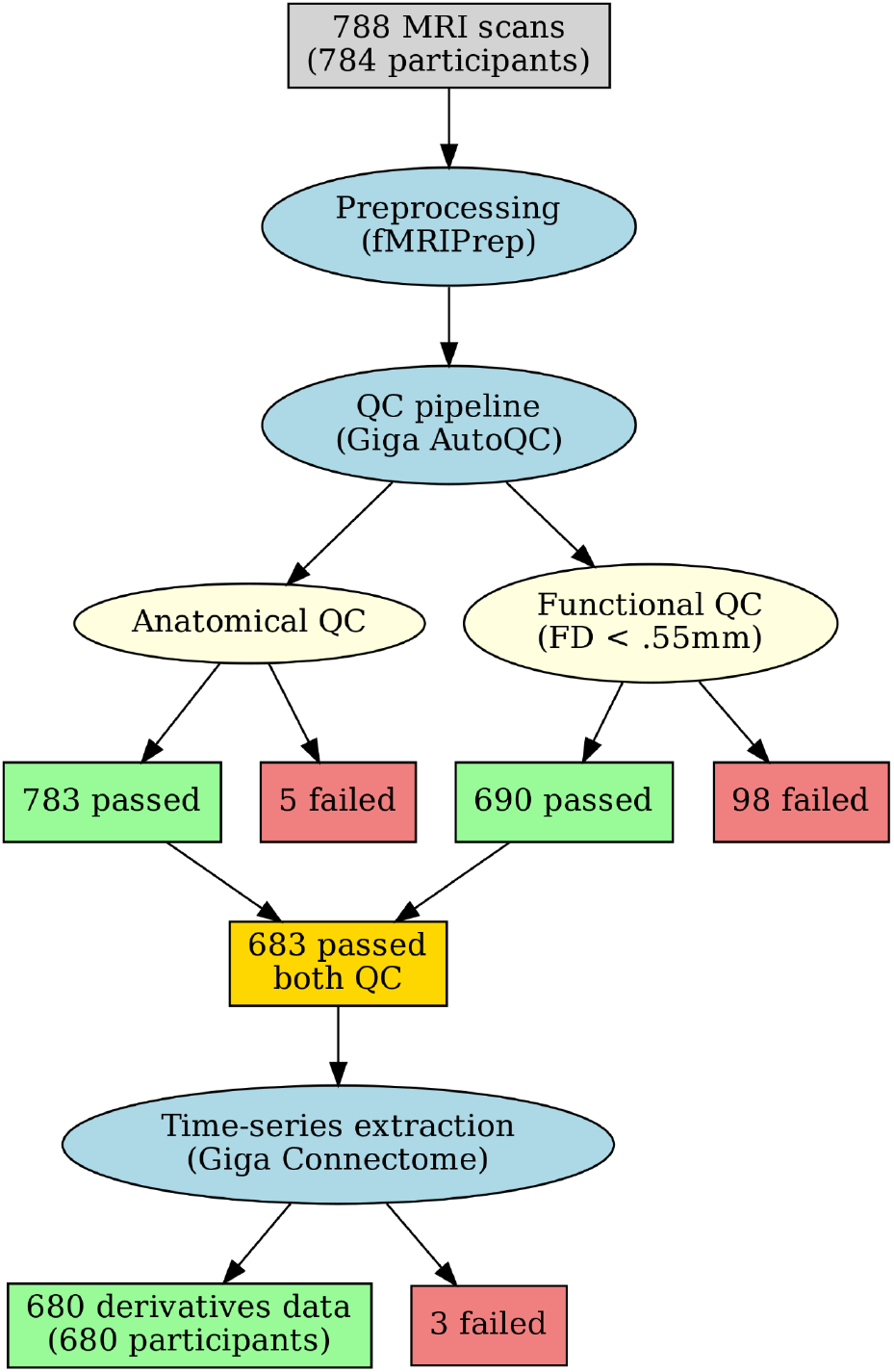
Flowchart showing steps and results of fMRI preprocessing, QC and time-series extraction.

Figure 2 shows the percentage of participants that passed QC within each diagnostic group. The distribution of mean FD by diagnostic group is shown in Figure 3. Across all participants there was a weak, positive correlation between motion and age (ρ = 0.1, p = .006) and a weak, negative correlation between motion and MoCA score (ρ = -0.12, p = .001), indicating that older individuals with greater cognitive impairment exhibited greater in-scanner movement (Figure 4). All data are available for download, including scans that passed or failed QC, with functional QC at both liberal and more stringent motion thresholds. We recommend that researchers with specific data quality requirements perform their own QC checks to ensure the data meets their standards.

**Figure 2.**
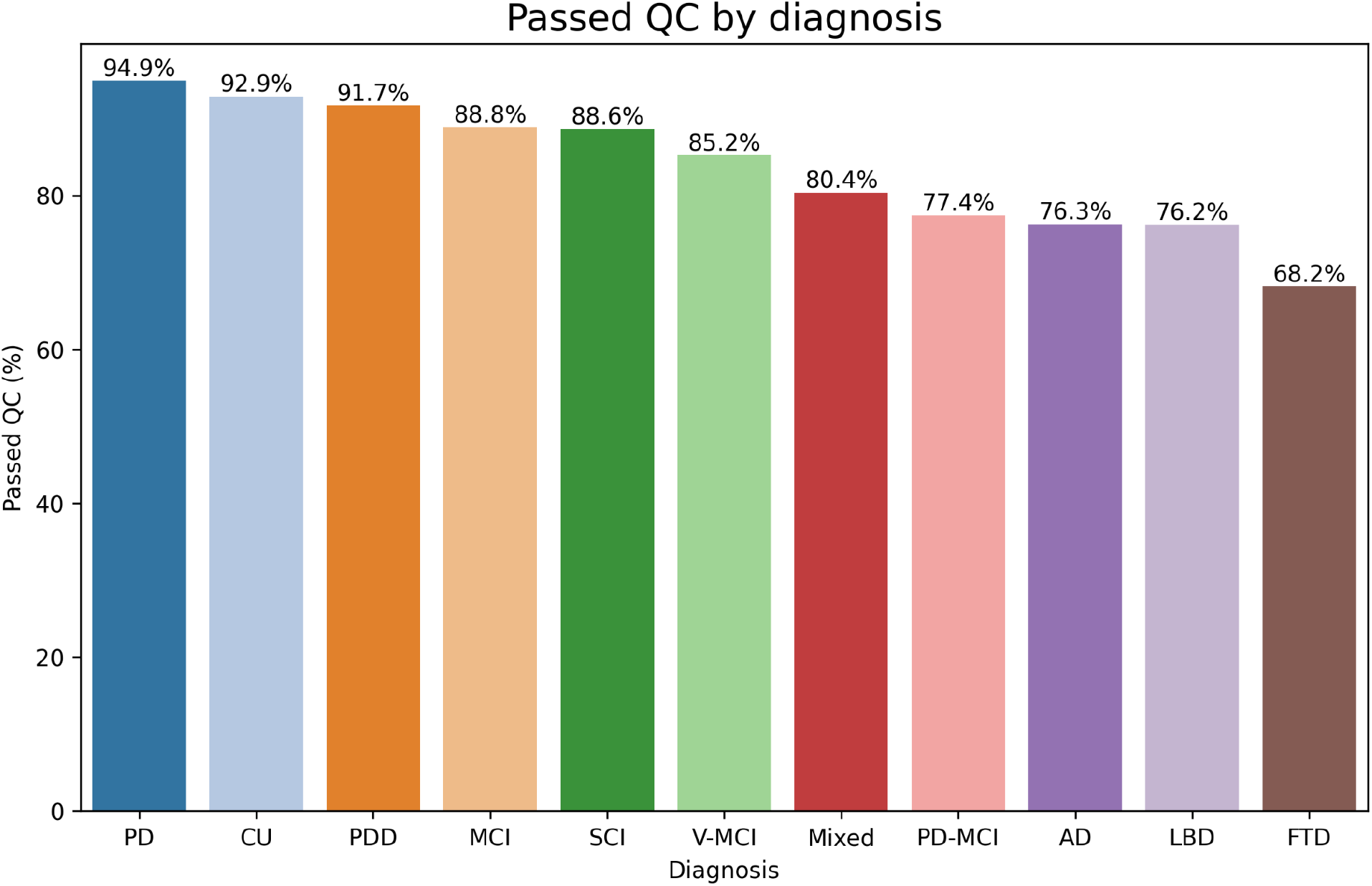
Percentage of participants that passed both anatomical and functional QC by diagnostic group. PD = Parkinson’s disease, CU = cognitively unimpaired, PDD = Parkinson’s disease with dementia, MCI = mild cognitive impairment, SCI = subjective cognitive impairment, V-MCI = vascular MCI, PD-MCI = Parkinson’s disease with MCI, AD = Alzheimer’s disease, LBD = Lewy body disease, FTD = frontotemporal dementia.

**Figure 3.**
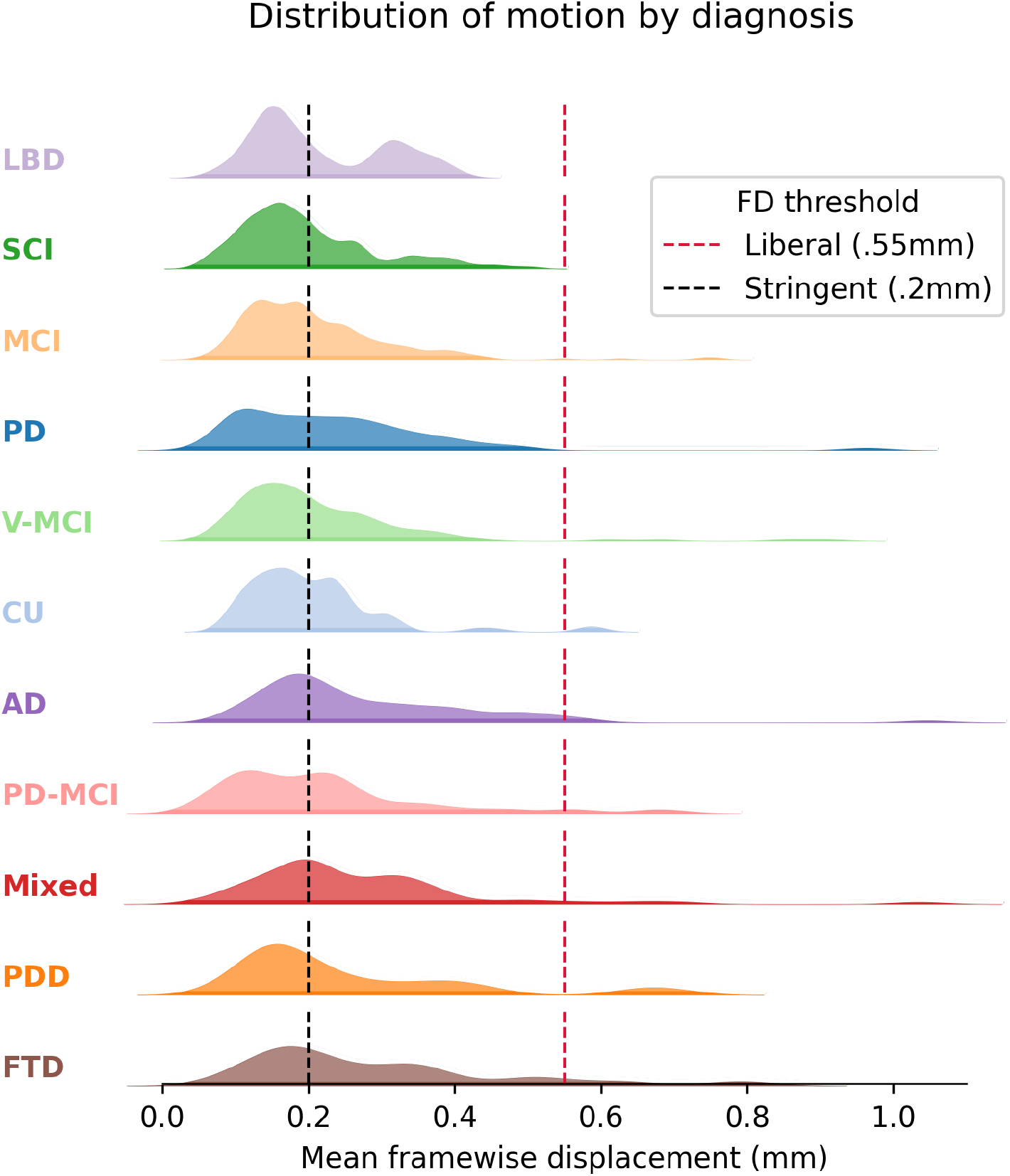
Distribution of mean framewide displacement (FD) by diagnostic group. Groups are ordered by the percentage of participants that fall under the liberal FD threshold. Note that one outlier with a mean FD of 17.4 is not shown. LBD = Lewy body disease, SCI = subjective cognitive impairment, MCI = mild cognitive impairment, PD = Parkinson’s disease, V-MCI = vascular MCI, CU = cognitively unimpaired, AD = Alzheimer’s disease, PD-MCI = Parkinson’s disease with MCI, PDD = Parkinson’s disease with dementia, FTD = frontotemporal dementia.

**Figure 4.**
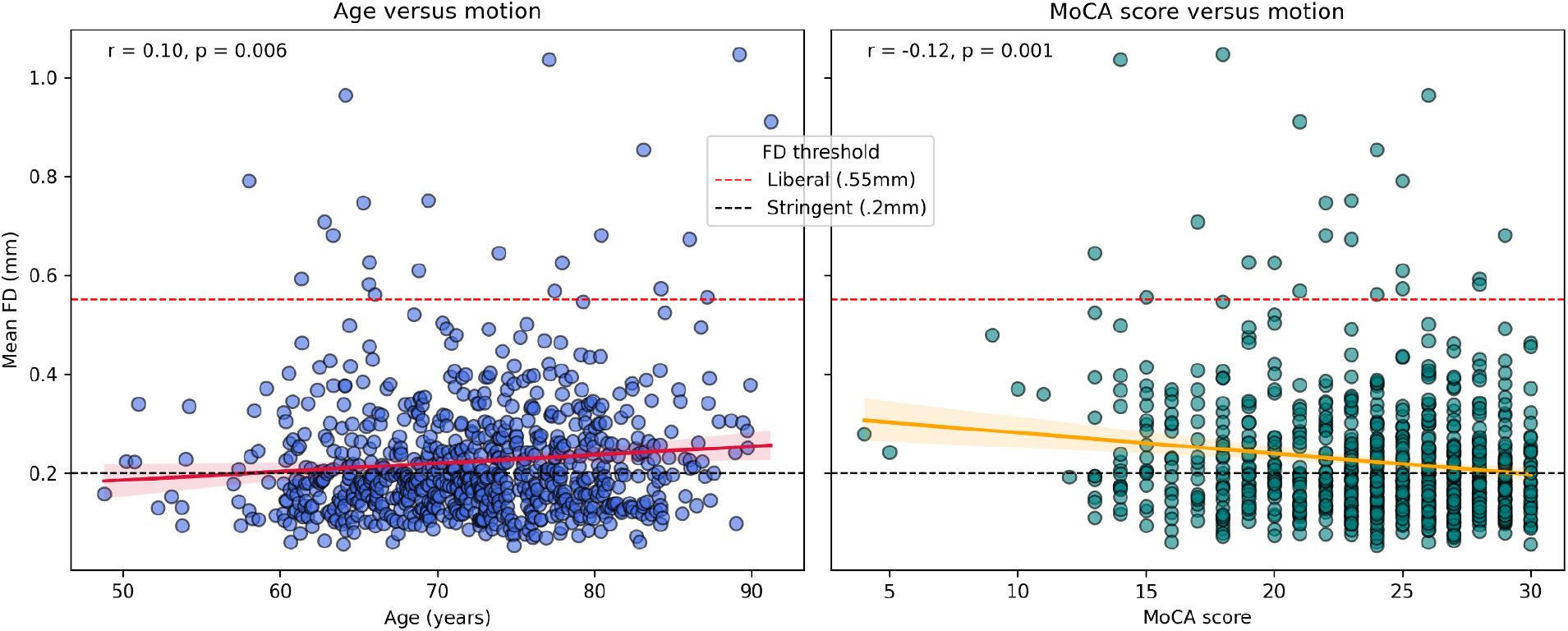
Relationship between mean framewise displacement (FD) and participant characteristics. Left: correlation with age; right: correlation with MoCA score. The MoCA is scored from 0 to 30, with higher scores indicating better cognitive function (note that one participant with an outlier score of 30.5 is not shown). Correlations were assessed using Spearman’s rho.

Overall, 86% of participants passed both anatomical and functional QC and had time-series derivatives available. The majority of QC failures were due to high motion, a common challenge in older populations with neurodegenerative conditions. This may introduce bias to the dataset, as participants who fail QC are likely to be older and more severely impaired (Figure 4). Motion and QC also varied across diagnostic categories (Figures 2 and 3), which may bias downstream analyses, for example if comparing different groups. Additionally, we did not investigate site effects, however, some sites are specialist centres focusing on recruiting particular diagnoses, which may introduce further bias. Nevertheless, the final dataset consists of a large number of high quality RSFC derivatives available to the research community, to further research on fMRI as a biomarker.

## Code availability

Code for the DICOM to BIDS conversion can be found at https://github.com/SIMEXP/bids_conversion. All MRI processing scripts are available on github: https://github.com/SIMEXP/giga_preprocess2/tree/compass_nd/compass-nd.

## Supplementary materials

### Anatomical preprocessing

Note that this text was automatically generated by fMRIPrep. A total of 1 T1-weighted (T1w) images were found within the input BIDS dataset.The T1-weighted (T1w) image was corrected for intensity non-uniformity (INU) with N4BiasFieldCorrection (Tustison et al. 2010), distributed with ANTs 2.3.3 (Avants et al. 2008, RRID:SCR_004757), and used as T1w-reference throughout the workflow. The T1w-reference was then skull-stripped with a *Nipype* implementation of the antsBrainExtraction.sh workflow (from ANTs), using OASIS30ANTs as target template. Brain tissue segmentation of cerebrospinal fluid (CSF), white-matter (WM) and gray-matter (GM) was performed on the brain-extracted T1w using fast (FSL 5.0.9, RRID:SCR_002823, Zhang, Brady, and Smith 2001). Brain surfaces were reconstructed using recon-all (FreeSurfer 6.0.1, RRID:SCR_001847, Dale, Fischl, and Sereno 1999), and the brain mask estimated previously was refined with a custom variation of the method to reconcile ANTs-derived and FreeSurfer-derived segmentations of the cortical gray-matter of Mindboggle (RRID:SCR_002438, Klein et al. 2017). Volume-based spatial normalization to two standard spaces (MNI152NLin2009cAsym, MNI152NLin6Asym) was performed through nonlinear registration with antsRegistration (ANTs 2.3.3), using brain-extracted versions of both T1w reference and the T1w template. The following templates were selected for spatial normalization: *ICBM 152 Nonlinear Asymmetrical template version 2009c* [Fonov et al. (2009), RRID:SCR_008796; TemplateFlow ID: MNI152NLin2009cAsym], *FSL’s MNI ICBM 152 non-linear 6th Generation Asymmetric Average Brain Stereotaxic Registration Model* [Evans et al. (2012), RRID:SCR_002823; TemplateFlow ID: MNI152NLin6Asym].

### Functional preprocessing

Note that this text was automatically generated by fMRIPrep. For each of the BOLD run found per subject (across all tasks and sessions), the following preprocessing was performed. First, a reference volume and its skull-stripped version were generated using a custom methodology of *fMRIPrep*. Susceptibility distortion correction (SDC) was omitted. The BOLD reference was then co-registered to the T1w reference using bbregister (FreeSurfer) which implements boundary-based registration (Greve and Fischl 2009). Co-registration was configured with six degrees of freedom. Head-motion parameters with respect to the BOLD reference (transformation matrices, and six corresponding rotation and translation parameters) are estimated before any spatiotemporal filtering using mcflirt (FSL 5.0.9, Jenkinson et al. 2002). The BOLD time-series (including slice-timing correction when applied) were resampled onto their original, native space by applying the transforms to correct for head-motion. These resampled BOLD time-series will be referred to as *preprocessed BOLD in original space*, or just *preprocessed BOLD*. The BOLD time-series were resampled into several standard spaces, correspondingly generating the following *spatially-normalized, preprocessed BOLD runs*: MNI152NLin2009cAsym, MNI152NLin6Asym. First, a reference volume and its skull-stripped version were generated using a custom methodology of *fMRIPrep*. Several confounding time-series were calculated based on the *preprocessed BOLD*: framewise displacement (FD), DVARS and three region-wise global signals. FD was computed using two formulations following Power (absolute sum of relative motions, Power et al. (2014)) and Jenkinson (relative root mean square displacement between affines, Jenkinson et al. (2002)). FD and DVARS are calculated for each functional run, both using their implementations in *Nipype* (following the definitions by Power et al. 2014). The three global signals are extracted within the CSF, the WM, and the whole-brain masks. Additionally, a set of physiological regressors were extracted to allow for component-based noise correction (*CompCor*, Behzadi et al. 2007). Principal components are estimated after high-pass filtering the *preprocessed BOLD* time-series (using a discrete cosine filter with 128s cut-off) for the two *CompCor* variants: temporal (tCompCor) and anatomical (aCompCor). tCompCor components are then calculated from the top 2% variable voxels within the brain mask. For aCompCor, three probabilistic masks (CSF, WM and combined CSF+WM) are generated in anatomical space. The implementation differs from that of Behzadi et al. in that instead of eroding the masks by 2 pixels on BOLD space, the aCompCor masks are subtracted a mask of pixels that likely contain a volume fraction of GM. This mask is obtained by dilating a GM mask extracted from the FreeSurfer’s *aseg* segmentation, and it ensures components are not extracted from voxels containing a minimal fraction of GM. Finally, these masks are resampled into BOLD space and binarized by thresholding at 0.99 (as in the original implementation). Components are also calculated separately within the WM and CSF masks. For each CompCor decomposition, the *k* components with the largest singular values are retained, such that the retained components’ time series are sufficient to explain 50 percent of variance across the nuisance mask (CSF, WM, combined, or temporal). The remaining components are dropped from consideration. The head-motion estimates calculated in the correction step were also placed within the corresponding confounds file. The confound time series derived from head motion estimates and global signals were expanded with the inclusion of temporal derivatives and quadratic terms for each (Satterthwaite et al. 2013). Frames that exceeded a threshold of 0.5 mm FD or 1.5 standardised DVARS were annotated as motion outliers. All resamplings can be performed with *a single interpolation step* by composing all the pertinent transformations (i.e. head-motion transform matrices, susceptibility distortion correction when available, and co-registrations to anatomical and output spaces). Gridded (volumetric) resamplings were performed using antsApplyTransforms (ANTs), configured with Lanczos interpolation to minimize the smoothing effects of other kernels (Lanczos 1964). Non-gridded (surface) resamplings were performed using mri_vol2surf (FreeSurfer).

